# Mechanisms and Safety of Ultrasound Phacoemulsification

**DOI:** 10.1101/2025.10.31.685733

**Authors:** Thomas Graf, Thomas Gisler, Silvio Emmenegger, Samuel Tanner, Rupert Menapace, Silvio Di Nardo

## Abstract

**Objective:** A comprehensive technical and biomedical analysis of ultrasound-induced effects during phacoemulsification cataract surgery (PCS) and an assessment of the potential risks associated with ultrasound emulsification

**Methods:** Modeling ultrasound wave propagation and intensities using the finite element method (FEM). Cavitation onset was experimentally validated in a pressure chamber observed with a high-speed camera. The thermal effects of ultrasound energy were estimated through established physical and thermal models and confirmed in a cuvette setup using thermistor probes.

**Results:** At the distal end of the PCS tip, the ultrasound wave intensity reached 0.2 W/mm^2^, resulting in ocular tissue displacements in the anterior chamber below 2 µm. The temperature rise in the anterior chamber was less than 0.2 °C/s. Both observations indicate negligible mechanical and thermal risk to ocular tissues. High-speed imaging confirmed cavitation confined to the tip region. Emulsification efficacy was maintained even under conditions that suppressed cavitation, including elevated ambient pressures and cavitation-inhibiting fluids. Results indicate that mechanical fragmentation by tip oscillation (“jackhammer effect”) is the only relevant emulsification mechanism, while cavitation plays a negligible role. Acoustic streaming was observed during ultrasound excitation without phaco sleeve. During phacoemulsification, the sleeve suppressed acoustic streaming into the anterior chamber.

**Conclusions:** The findings validate the safety of ultrasound-based PCS and confirm that tissue fragmentation is primarily driven by the oscillating tip’s direct action.

## Introduction

Today, two main technologies to workup the cataractous lens are available: ultrasound-based phacoemulsification cataract surgery (PCS) and femtolaser assisted cataract surgery (FLACS). Since the introduction of FLACS in 2011, numerous randomized controlled trials[1–3] and systematic reviews[4–6] to compare safety and effectiveness of the two technologies have been published. Considering the current knowledge, the American Academy of Ophthalmology (AAO) concludes that FLACS and PCS have similar excellent safety and refractive outcomes but that FLACS is less cost-effective[7]. The European Society of Cataract & Refractive Surgeons (ESCRS) has published a similar positioning in a draft of a guideline[8]. Thus, from the clinical perspective, ultrasound-based PCS is a well-proven and safe technology, enabling surgeons to achieve the best-possible clinical results. Ultrasound will remain the most important and widely used technology for cataract removal. It is therefore important to continuously improve safety and effectiveness of this method.

The interaction between ultrasound and proteins is mediated by six physico-chemical effects: a) ultrasound pressure, b) cavitation c) acoustic streaming, d) temperature e) free radicals and f) shear forces[9]. The aim of this study is to analyze and evaluate these physico-chemical effects by numerical simulation and by experiment using a modern, state-of-the-art phacoemulsification system.

The results concerning free radical generation have been published elsewhere[10]: experiments demonstrated that free radicals are generated during the delivery of ultrasonic power. Air-bubbles in the fluid surrounding the PCS tip enhance the generation of free radicals, whereas running fluidics and the emulsification process itself of lens tissue reduce the production of free radicals. In the clinically desired situation, the PCS tip is occluded by the lens and only as much ultrasound energy as needed is applied; in this situation, a small peak in the Electron-Paramagnetic Resonance (EPR) spectrum representing the number of produced radicals is visible. Due to the minimal intensity of the peak, the corresponding number of free radicals lies below the limit of accurate quantification and can therefore be considered negligible. The authors conclude that other causes than ultrasound might be responsible for suggested oxidative stress during cataract surgery.

In this work, five ultrasound-specific, safety-relevant processes were investigated through technical analysis: propagation of ultrasound waves in the anterior chamber, formation and dynamics of cavitation bubbles, generation of acoustic streaming, the generation of shear forces and the temperature rise in the anterior chamber. Up to now, no systematic assessment of all the relevant ultrasound-induced risks on an engineering level has been published.

## Methods

### Numerical Simulations

Numerical simulations were performed in COMSOL Multiphysics® (v.6.1 & 6.2. COMSOL AB, Stockholm, Sweden) using the finite element method (FEM). Coupled wave propagation in both water and solid materials was computed in frequency-domain studies. A 3D model of the human eye and the PCS tip and handpiece was created in the software. The longitudinal oscillation of the tip was driven by a prescribed displacement of the piezoelectric package operating at 27 kHz. The resulting motion of the tip and shaft agrees to within 5 µm with measurements obtained using a laser Doppler vibrometer (Polytec; Controller: OFV-3320, Sensor: LSV-065-306F), with a maximum tip amplitude of 50 µm. The elastic modulus, Poisson’s ratio, and mass density of titanium (Grade 5) for the PCS tip and handpiece, as well as those of the elastomer (Elastosil® LR 3003/85) for the sleeve, were defined as material properties in the simulation. Realistic viscoelastic models and material properties for ocular tissues were obtained from the literature and integrated into the simulation, including distinct internal damping models for the cornea[11–13]. The fluid in the anterior chamber of the eye was modeled as water. From all the materials, the elastomer exhibited the lowest damping, with a Q-factor of approximately 20, corresponding to a characteristic relaxation time of *τ* = 0.2 ms at a frequency of 27 kHz. As this relaxation (or transient) time is significantly shorter than the phacoemulsification application duration, frequency-domain studies are appropriate for FEM simulations.

Supplementary simulations were conducted using the Heat Transfer module of COMSOL Multiphysics®. These simulations investigated the heat generation resulting from the movement of the tip in a viscous liquid.

### Experimental methods

#### Phacoemulsification Apparatus with Handpiece

Phacoemulsification was performed using two surgical platforms: the CataRhex 3 easy and the OS4 (both manufactured by Oertli Instrumente AG, Berneck, Switzerland). The procedure utilized the easyPhaco handpiece (Oertli), which incorporates six piezoelectric ceramic rings. A variety of phaco tips (easyTip, Oertli) were employed, including CO-MICS tips (1.6 mm, angled at 30° and 53°) and standard tips (2.2 mm and 2.8 mm, angled at 15°, 30°, and 40°). All tips were manufactured from titanium grade 5. The handpiece was operated in continuous ultrasound mode to deliver a constant energy output throughout the procedure.

#### High-Speed Camera and Pressure Chamber

Cavitation bubbles were observed using a high-speed camera (Motion Pro X4, Redlake) and macro lenses. Up to seven images could be captured during a single oscillation cycle of 37 µs at 27 kHz. To suppress the formation of cavitation bubbles during ultrasonic excitation, a pressure chamber capable of reaching 4 bar ambient pressure was constructed, see Fig. 1. The design concept was inspired by the work of J. Zacharias[14, 15]. A feedthrough for the PCS handpiece was incorporated, allowing phaco operation inside the chamber at a maximum ambient pressure of 4 bar. Through glass windows in the pressure chamber, the formation and dissipation of cavitation bubbles during phaco operation could be observed using a high-speed camera and intense LED illumination.

**Fig. 1.**
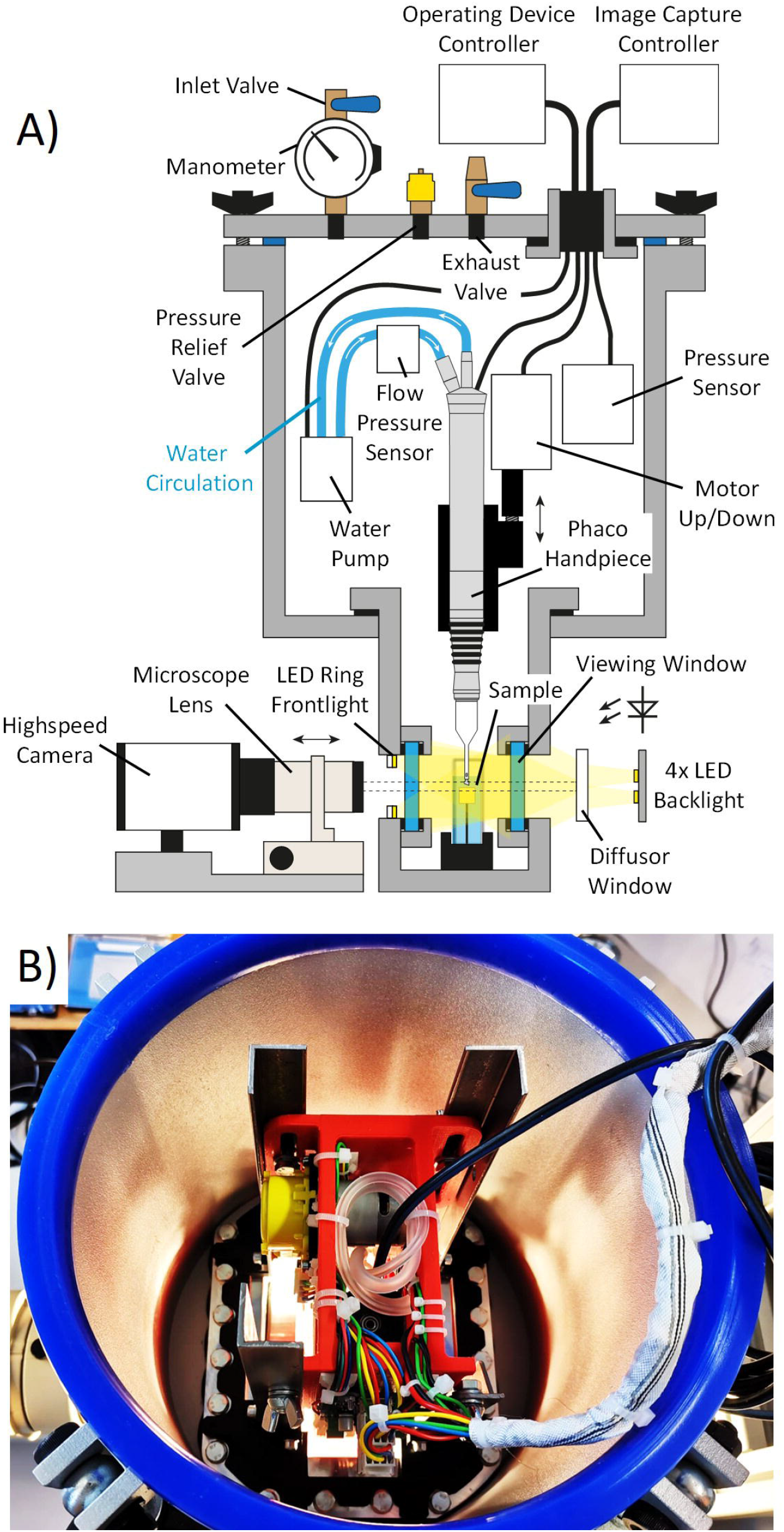
Pressure chamber. Pressure chamber for phacoemulsification under high ambient pressure. A) Schematic of the pressure chamber setup. The PCS handpiece can be positioned near the probe to enable phacoemulsification under elevated pressure. High-intensity LED illumination is used to facilitate visualization with the high-speed camera in backlight. Microscope lenses allow detailed observation of the probe and tip region. B) Photograph showing components within the chamber. The inner diameter of the chamber is 20.5 cm.

#### Phaco Emulsification Efficiency Measurement

Phaco times were measured at ambient pressures of 1 bar and 4 bar for the following ultrasound power settings: 20 %, 40 %, 60 %, 80 % and 100 %. An EasyTip 2.2 mm 40° phaco tip attached to an OS4 unit operating in continuous mode served as the emulsification instrument. The test substrates comprised 1 cm^3^ cubes of Sbrinz cheese (aged 22 months) and hardened pig lenses (4% formalin for 2 hours). Upon occurrence of occlusion, timing commenced; measurement ceased when a visible fluid jet emerged from the opposite face of the test substrates. At least 9 replicates were performed for each condition.

#### Temperature Measurement in a Cuvette Setup

Heat generation in water due to ultrasonic motion of the PCS tip was investigated using the experimental setup shown in the photograph of Fig. 2. The PCS tip was immersed in thermally insulated cuvettes containing either 4 g or 64 g of water. Water temperature was monitored using a thermistor probe (Fluke, 5640 series).

**Fig. 2.**
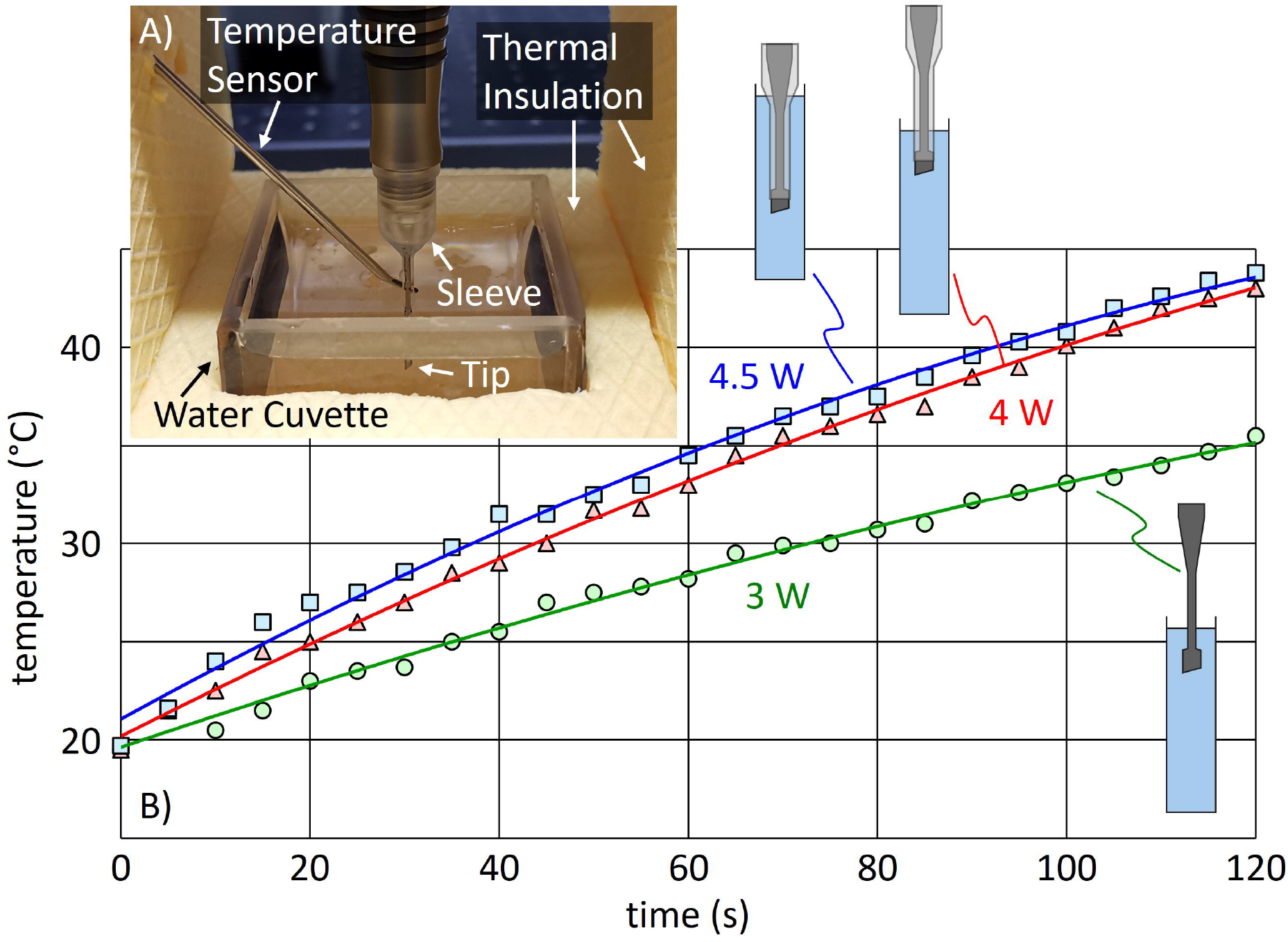
Heat generation. Heating power during phaco operation. A) The photograph shows the experimental setup. B) The graph indicates the temperature rise of 4 g water as a function of time with different operation modes: fully immersed with sleeve (squares), partially immersed with sleeve (triangles) and partially immersed without sleeve (circles). The three curves allow to calculate the heating power (see text).

#### Infrared Camera

Heat generation of the PCS handpiece was investigated using an infrared camera (Pyroview from Infrared Systems). To allow for direct view of the heating of the piezoelectric components and wiring, the casing of the PCS handpiece was removed. Heat-resistant black paint was applied to increase the emissivity of the metal parts to close to 1, ensuring accurate temperature measurements.

## Results

### Simulation Results of Ultrasound and Cavitation

Oscillation of the PCS tip and the metal shaft of the handpiece generates ultrasound waves in the fluid of the anterior chamber, as well as in the irrigation and aspiration channels within the handpiece. Fig. 3 illustrates the simulated acoustic pressure amplitudes in the central plane through the anterior chamber of the eye, encompassing the PCS tip and the distal segment of the handpiece. Acoustic pressures of up to 5 bar are observed directly in front of the PCS tip and within its distal portion. Acoustic pressure exhibits rapid attenuation in the corneal direction, reaching 0.1 bar at 3 mm from the distal center of the tip. The acoustic pressure of the fluid within the anterior chamber induces displacements in the viscoelastic tissues of the eye. These displacements remain well below 2 µm across all ocular structures, notably in the corneal region.

**Fig. 3.**
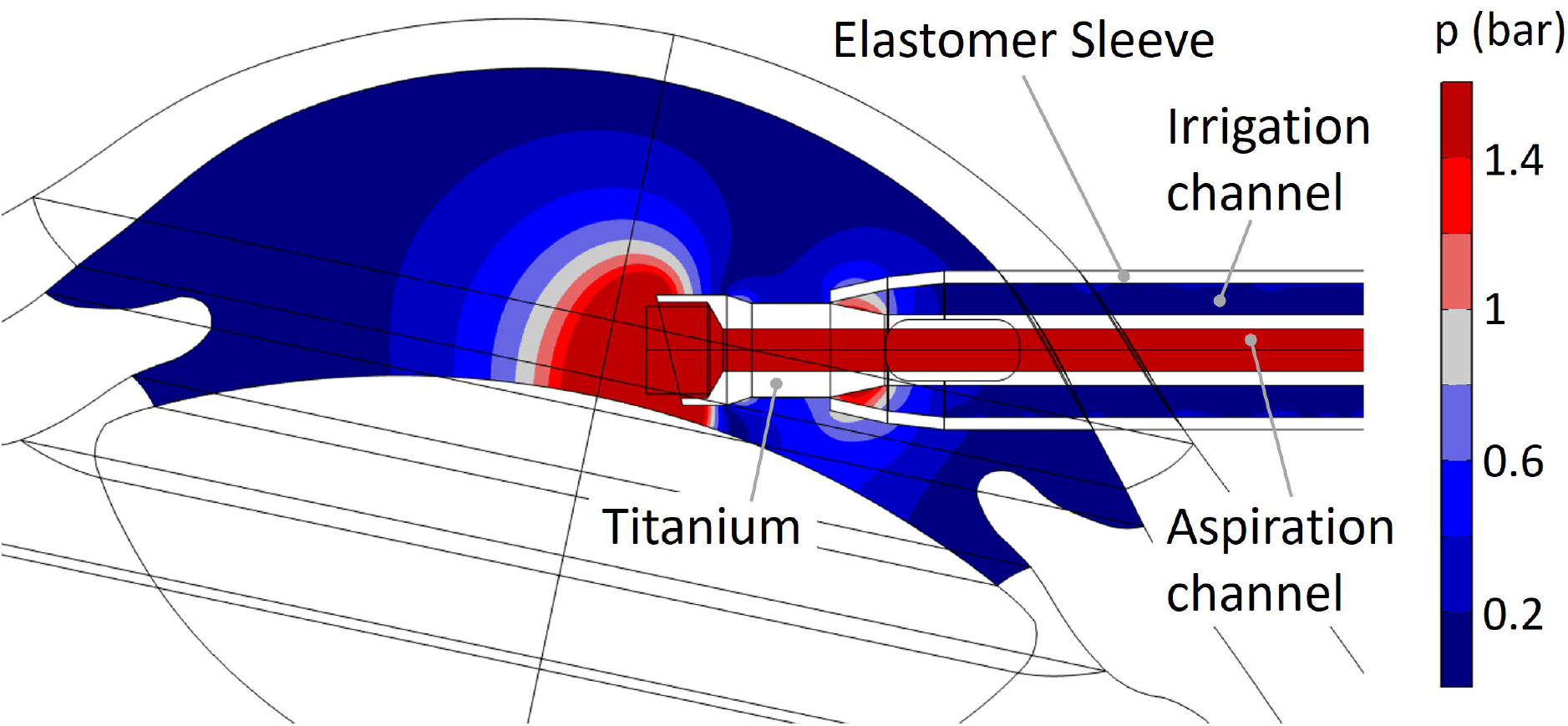
Acoustic pressure. Simulated acoustic pressure amplitudes in the aqueous humor of the anterior chamber and at the distal end of the PCS tip. Cavitation bubble formation is possible in the red regions, where the acoustic pressure amplitude exceeds 1 bar.

### Experimental Results

#### Observation of Cavitation Vapor Bubbles and Their Suppression

To rule out the potential formation of air bubbles, we used boiled and distilled water for our experiments. Where the simulated acoustic pressure locally exceeds 1 bar (Fig. 3), cavitation, i.e. the formation of vapor bubbles during the negative half-cycle becomes possible. Fig. 4.A) shows a high-speed photograph of the PCS tip shaft, revealing cavitation bubbles forming at locations predicted by simulations (not visible in the section shown in Fig. 3). The graph of Fig. 4.B) displays the observed bubble area within the section of Fig 4.A) as a function of observational time. The measured bubble area is best fit by a sine curve at 27.1 kHz, indicating that the bubbles pulsate at the tip’s expected oscillation frequency and that cavitation occurs where predicted by the simulation. The number of vapor bubbles is significantly lower than anticipated given the extensive regions of high ultrasound pressure in the simulations. This discrepancy may be due to the lack of nucleation sites in the pure water used.

**Fig. 4.**
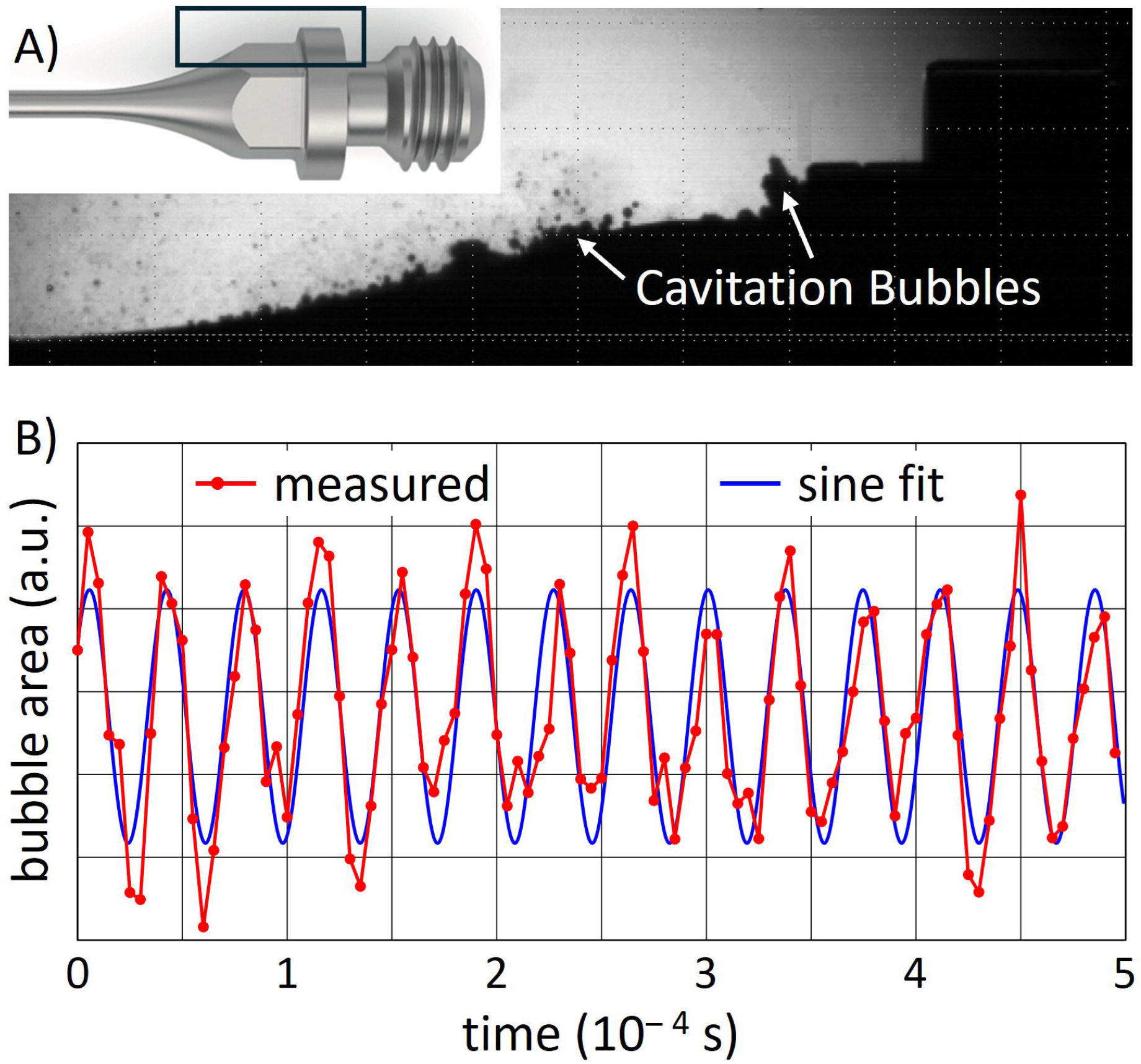
Formation of cavitation bubbles. Observation of cavitation bubbles. A) Image from a high-speed sequence showing cavitation bubbles on the shaft of the PCS tip, without the sleeve. The tip oscillates at full phaco power, and water circulation is disabled. The rectangle of the inset depicts the region of observation. B) The graph shows the area of the cavitation bubbles as a function of time, measured every 5 µs (red dots). The fitted blue curve shows the cavitation bubble pulsation at 27.1 kHz.

We observed cavitation at the shaft and the tip of the PCS handpiece, but never off the handpiece parts. This finding is confirmed by published photographs in the literature[14, 16]. The formation of cavitation bubbles in front and inside the tip supports the hypothesis that phaco emulsification may occur through the implosion of cavitation bubbles. Whether the observed cavitation bubbles at the distal end of the tip and inside the tip actually contribute to the emulsification of the cataractous lens will be clarified in the next chapter. FEM simulations of Fig. 3. show that the acoustic pressure is highest within the aspiration channel of the PCS tip, making cavitation and formation of vapor bubbles most likely in this region. To investigate local density variations between pure water and vapor-like regions containing cavitation bubbles, X-ray radiographic measurements were conducted using a d2 system from Diondo (Hattingen, Germany). However, the X-ray imaging technique was unable to resolve the density contrast between the two water phases, primarily due to attenuation and masking effects caused by the metal components of the tip.

In a separate experiment, the PCS tip was excited at 27 kHz within the pressure chamber under varying ambient pressures, up to a maximum of 4 bar. To increase the potential formation of cavitation bubbles in front of the tip, the conical section of the tip was cut as indicated in Fig. 5. In a separate experiment, the tip was cut perpendicularly further back, at the narrow section of the shaft. In all experiments, no cavitation was observed either in front of or within the conical section of the tip at ambient pressures of 3 bar or higher. An identical threshold pressure was reported by Zacharias (2008)[14].

**Fig. 5.**
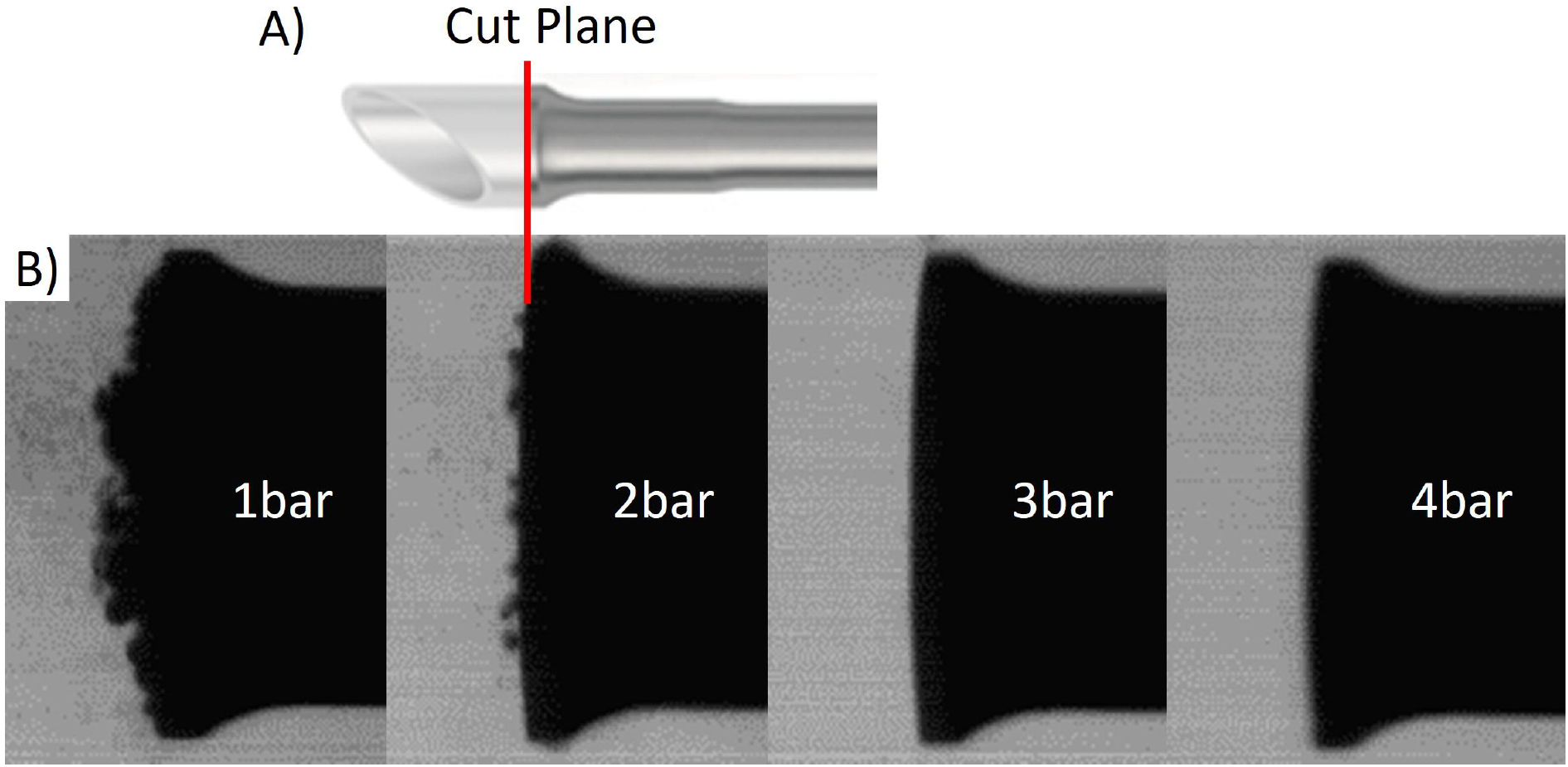
Pressure dependent cavitation formation. High-speed imaging of cavitation bubble activity at the PCS tip under varying ambient pressures inside the pressure chamber. A) The PCS tip was modified by cutting, to enhance cavitation bubble generation during oscillation. B) The tip oscillates at 27 kHz under full phaco power, with water circulation disabled. The image sequence captures oscillation events at ambient pressures ranging from 1 to 4 bar. Cavitation bubbles are clearly visible at lower pressures but are no longer observed at pressures of 3 bar and above.

#### Phaco Emulsification Efficiency

Following the assessment of cavitation bubble formation at ambient pressures up to 4 bar, the phacoemulsification process was subsequently investigated within the high-pressure chamber. The tip was brought into contact with a model tissue consisting of cubic samples (1 cm^3^) of hard Sbrinz cheese and phacoemulsification was initiated. The cheese cubes exhibited consistent material properties, which allowed for reproducible and comparable experimental conditions across different tests. Emulsification efficiency was evaluated by measuring the time t_p_ required for the PCS tip to completely pierce 1 cm through the model tissue. Repeated experiments showed that emulsification efficiency did not decrease at ambient pressures above 3 bar, where cavitation in front of or within the tip was suppressed.

The same experiments were also performed on porcine eye lenses that had been hardened by immersion for 2 hours in 4% formalin. In these specimens as well, phacoemulsification efficiency did not decrease with elevated ambient pressure.

In additional experiments, glycerin was used as the working fluid instead of water. Glycerin has a much larger critical radius than water, beyond which vapor pressure can overcome surface tension and allow a vapor bubble to grow freely. As a result, cavitation bubble formation in glycerin was absent. On the other hand, occlusion with glycerin was very weak. Nevertheless, phacoemulsification of the model tissue was still effective with glycerin, with a piercing time of approximately t_p_ = 6 seconds.

In a third experiment, a plant leaf was used to investigate the interaction of cavitation bubbles with organic tissue. In Fig. 6A, the tip of the leaf was placed in direct contact with the metal components within the conical section of the PCS tip. After 120 seconds of phacoemulsification in water, the leaf showed clear signs of wear, attributable to mechanical friction with the metal parts of the tip shaft. In contrast, in Fig. 6B, the leaf was positioned without any contact with the metal components of the PCS tip. After 120 seconds of phaco tip oscillation in water, no visible damage to the leaf was observed, despite the high likelihood of cavitation implosions occurring near the tissue.

**Fig. 6.**
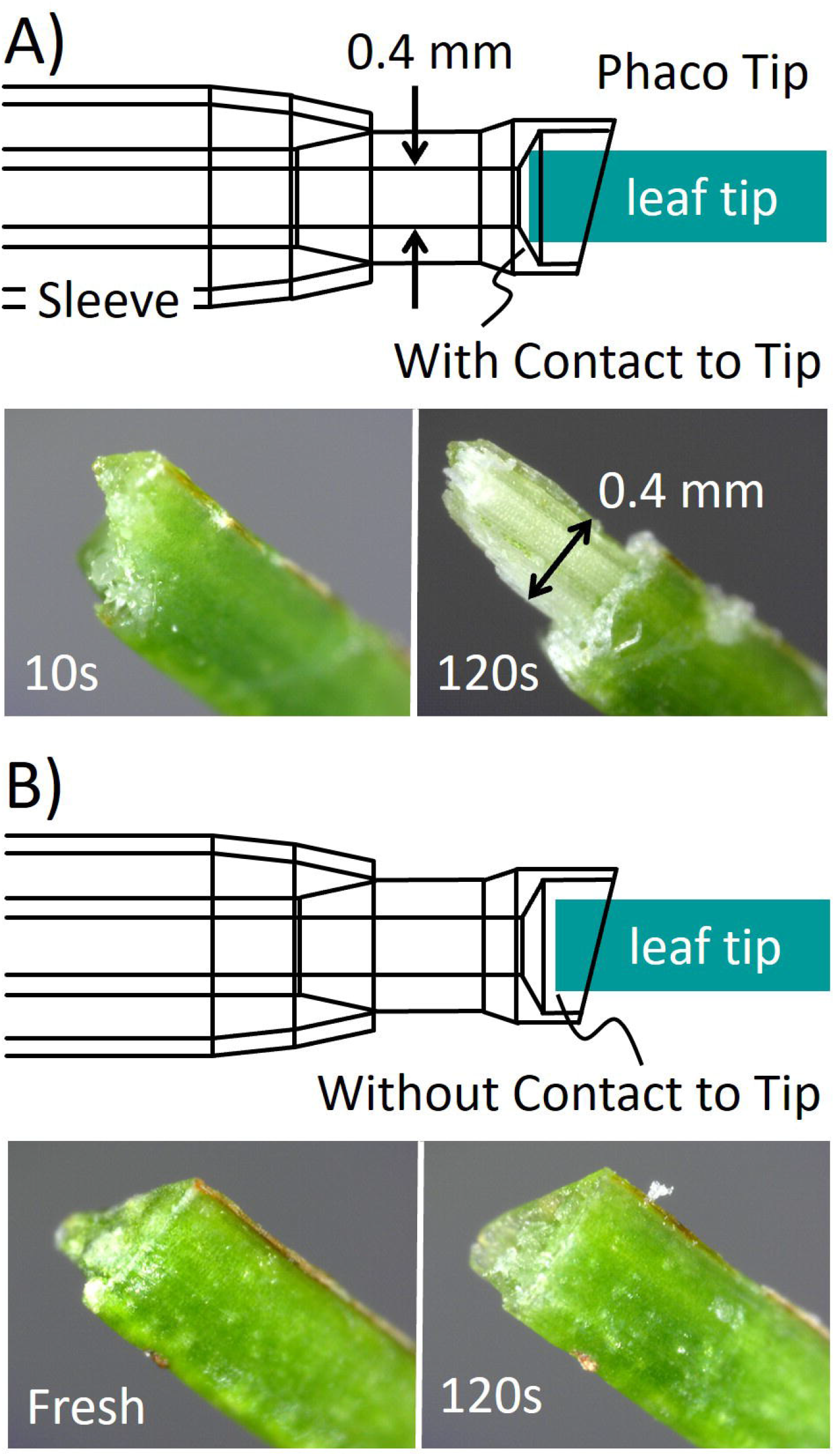
Interaction of cavitation bubbles with organic tissue. Phacoemulsification of model tissue in water. A leaf tip is inserted into the PCS tip. The photographs show removal (ablation) of the leaf tissue during phaco operation with mechanical contact between the leaf and the tip (A) and no significant removal without contact (B).

All the above experimental observations support the conclusion that cavitation is not the primary mechanism driving the emulsification of organic material during PCS.

#### Acoustic Streaming by Tip Oscillations

High-frequency oscillation of the PCS tip generates acoustic streaming, which primarily occurs when the sleeve is removed, exposing the vertical surface where the tip connects to the handpiece (see highlighted area in Fig. 4A). Video analysis of the tip oscillation in water estimated the resulting streaming velocity of cavitation bubbles to be approximately 0.1 m/s. By comparison, this is lower than the irrigation flow velocity observed from the sleeve’s infusion ports[15] and two orders of magnitude lower than the tip’s peak oscillation velocity of 8.5 m/s at 27 kHz with a 50 μm amplitude. The sleeve effectively suppresses acoustic streaming into the anterior chamber during phacoemulsification. Weaker streaming is also observed at the distal end of the tip. When aspiration is turned off, this streaming pushes particles in front of the tip away from it. Given the small oscillating vertical surface, the distal end of tip transfers minimal momentum to the fluid, resulting in limited ultrasound-driven streaming. This opposing flow is effectively overcome by aspiration through the tip, which is essential for stabilizing tissue fragments and ensuring efficient emulsification.

#### Heat Generation in the Anterior Chamber of the Eye

The heat generated by the oscillatory motion of the PCS tip within the anterior chamber of the eye can be estimated analytically. The tip is modeled as a cylinder with an outer diameter of D = 0.74 mm and length L = 2 cm. The surrounding water is assumed to follow the oscillatory motion of the cylinder surface, with the velocity field decaying exponentially with the radial distance y from the cylinder surface, described by:

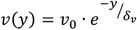

Here, 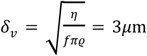 is the characteristic boundary layer thickness, and 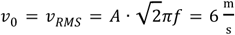 represents the root-mean-square velocity of oscillation. The viscosity and density of water at 36°C are taken as *η* = 7 × 10^−4^Pa · s and 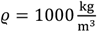, respectively. FEM simulations, incorporating a no-slip boundary condition at the water–cylinder interface, confirmed the exponential velocity profile and the boundary layer thickness. The viscous frictional force exerted by the fluid on the cylinder is calculated as:

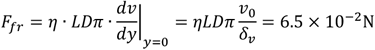

The corresponding heating power due to this frictional force is:

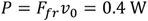

Accounting for both the inner and outer surfaces of the tip shaft, and considering that only the distal portion of the tip is immersed in the ocular fluid, the total heating power generated by the oscillating PCS tip in water remains well below 1 W.

The heat input into the anterior chamber of the eye was emulated through controlled experiments. In these experiments, the PCS tip was immersed to varying depths in thermally insulated cuvettes containing either 4 mL or 64 mL of water (see Fig. 2). The water temperature inside the cuvettes was recorded over a 2-minute interval under continuous application of maximum phaco power. The time-dependent temperature rise *T*(*t*) was observed to follow an exponential behavior:

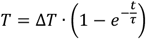

where Δ*T* and *τ* are fitting parameters derived from the experimental data. From this, the total heat power P delivered by the PCS tip to the water was calculated using:

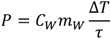

Here, 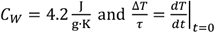 is the specific heat capacity of water, *m*_*W*_ is the mass of the water, and 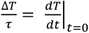 represents the initial rate of temperature increase.

Across multiple measurements, the estimated heat input was approximately 4 W. This value is an order of magnitude higher than the estimate based solely on viscous friction between the oscillating tip and the surrounding fluid, as discussed earlier. The discrepancy may be attributed to additional lateral motion of the tip within the fluid, friction between the tip and the sleeve, and more significantly, to heat conduction from the piezo actuator inside the PCS handpiece.

When extrapolated to the entire eye, which contains approximately 6 g of aqueous fluid, a heating power of 4 W corresponds to an initial temperature rise of:

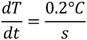

It is important to note that this temperature increase was observed under thermally insulated, stagnant fluid conditions.

The typical mean Overall Effective Phacotime is 10.72s for low fluidic settings and 7.44s for high fluidic settings[17]. This corresponds to a hypothetical maximum temperature rise of 2.1°C. As during the procedure, periods of ultrasound delivery alternate with phases of fragment removal by fluidics, the effective temperature rise is considerably lower than the hypothetical temperature rise, due to the cooling effect of the fluidics Thermal imaging using an infrared camera revealed that, in air and with the open handpiece, the temperature of the six piezoelectric rings increased by several degrees within the first few seconds of activation. In addition to the piezoelectric elements, the piezo driver circuit contributes to heat generation. This heat is transferred via the irrigation fluid into the anterior chamber of the eye, but is removed during periods of an unoccluded tip through aspiration via the handpiece. The volume of fluid transported corresponds to a thermal transport capacity of 150 J/(K·min), which appears sufficient to outbalance any clinically significant temperature rise.

## Discussion

Numerical simulations and experimental investigations consistently demonstrated the following:

### Emulsification Mechanism: Jackhammer, Not Cavitation

- **Cavitation Presence and Impact**: Numerical simulations confirmed that cavitation bubbles formed only at the PCS tip and shaft, restricted to zones of high acoustic pressure. Cavitation in front of the tip at ambient pressures ≥3 bar was suppressed. The absence of cavitation did not reduce emulsification efficiency, suggesting that cavitation is not essential for effective tissue fragmentation.
- **Medium Comparison**: In glycerin, which strongly inhibits cavitation, phacoemulsification was still effective, albeit about 50% slower, partly due to reduced occlusion efficiency. This further supports a mechanical, rather than cavitation-based, emulsification process.
- **Mechanical Contact Experiments**: Experiments with plant tissue confirmed that cavitation alone, in the absence of direct mechanical contact, caused no observable damage. Tissue wear was only evident where physical contact with oscillating metal surfaces occurred.
- **Proposed Mechanism of Phacoemulsification:** Lens fragments are stabilized at the distal end of the PCS tip by aspiration vacuum and their own inertia, allowing mechanical disintegration by the longitudinally oscillating beveled tip (jackhammer effect). The fragments are then drawn into the aspiration channel, where high-frequency wall motion and fragment inertia prevent clogging and promote further disintegration through frictional interaction with the channel walls. Ultrasound frequency is crucial for generating high accelerations that leverage fragment inertia. The formation of cavitation bubbles is a secondary effect, with vapor bubble collapses being infrequent and not essential to the emulsification mechanism.

### Minimal Risk to Corneal and Intraocular Tissues

- **Acoustic Pressure and Displacement**: Acoustic pressures from PCS peaked at 5 bar near the tip and declined to 0.1 bar at the inner surface of the cornea, producing tissue displacements below 2 µm— significantly lower than those caused by manual surgical handling.
- **Temperature Rise**: Heat generated by PCS tip oscillation was analytically and experimentally determined to be <4 W, leading to an estimated temperature increase of <0.2°C/s in the anterior chamber. Typically, the temperature increase is below 2°C during phacoemulsification. In comparison, the temperature rise of the cornea during excimer laser refractive surgery has been measured using two different surgical platforms and are 7.2°C and 4.1°C respectively[18]. Thermal damage in the cornea is expected to be observed after a heat dose of 45°C during 45 minutes[19]. In another study, a thermal impact of 40°C for 10sec did not show histological changes in the cornea[20]. Thus, thermal damage due to the thermal impact of the ultrasound can be excluded.
- **Streaming and Flow Effects**: Ultrasound-induced acoustic streaming is effectively suppressed by the sleeve and by the aspiration flow when active. Consequently, it does not impair emulsification efficiency or pose a risk to ocular tissue.

## Conclusion

This study provides strong evidence that ultrasound phacoemulsification using a PCS handpiece functions primarily through a **mechanical jackhammer mechanism**, and not by cavitation. Both simulation and experimental data confirm that the system **does not pose a thermal or mechanical risk** to sensitive ocular tissues, including the cornea. The observed effects—limited sound pressure propagation, negligible tissue displacement, and minor temperature increase—support the safety and efficacy of this technology for clinical ophthalmic use.

## Abbreviations

AAO: American Academy of Ophthalmology
EPR: Electron-Paramagnetic Resonance
ESCRS: European Society of Cataract & Refractive Surgeons
FEM: Finite Element Method
FLACS: Femtolaser Assisted Cataract Surgery
PCS: Phacoemulsification Cataract Surgery

